# Drivers of phenotypic differentiation across rock outcrop sky islands in *Impatiens* plants

**DOI:** 10.1101/2023.11.01.565130

**Authors:** Sumayya Abdul Rahim, Aboli Kulkarni, Ullasa Kodandaramaiah

## Abstract

Habitat specialization can lead to naturally fragmented populations separated by unoccupied habitats that comprise potential barriers to dispersal. Habitat fragmentation can result in phenotypic divergence between populations, and eventual speciation. We investigate trait divergence in a habitat specialist plant species – *Impatiens lawii* – that is specialized on rock outcrops in the Western Ghats mountains of India. We compare trait divergence in this species with that in a closely related species, *I. oppositifolia*, which is a continuously distributed habitat generalist that co-occurs with the former species on rock outcrops in addition to being found in intervening regions. As predicted, we found that floral traits were more strongly structured across populations in *I. lawii* than in *I. oppositifolia*, supporting the role of habitat specialization. In both species, floral traits were more strongly structured across populations than were vegetative traits. Interestingly, the population pairs that were significantly differentiated for floral traits differed from those differentiated for vegetative traits. Furthermore, some population pairs differed in vegetative traits but not in floral traits. The incongruence in patterns of floral and vegetative trait differentiation suggests the role of selection. There was no association between floral trait variation and geographic distance in either species. Surprisingly, there was a significant correlation between geographic distance and vegetative trait variation in *I. oppositifolia*, but not in *I. lawii* (Fig 4D; Table 2), possibly due to genetic drift. Overall, a combination of habitat specificity, drift and selection best explains trait variation in *I. lawii*.

## Introduction

The influence of niche specialization on diversification has interested many researchers and spanned a multitude of studies across taxa. Transitions from generalization to specialization have occurred repeatedly across the tree of life (Vamosi et al. 2014). Studies investigating multiple axes of specialization have reported both increase (e.g.,Salisbury et al. 2012; Hardy and Otto 2014) and decrease (e.g., Fernández-Mazuecos et al. 2013; Cyriac and Kodandaramaiah 2018) in diversification rates of lineages following a switch to specialization. Therefore, niche specialization can have significant consequences on speciation and diversification of taxa, but these effects are not well understood. Habitat specialization is one axis of speciation wherein species occupy subsets of habitats available to them. In species with multiple populations, habitat specialization leads to naturally fragmented populations separated by other habitats that comprise potential barriers to dispersal. Habitat fragmentation associated with habitat specialization can increase the probability of extinction (Colles et al. 2009; Püttker et al. 2013). On the other hand, habitat specialization can also lead to increased diversification by promoting divergences among populations (Rice 1987; Sacks et al. 2008; Lenormand 2012; Berner and Thibert-Plante 2015).

Sky islands present excellent study systems to understand the effects of habitat specialization on population divergence and intra-specific diversification. Sky islands are high altitude, fragmented, montane habitats that are separated by valleys with distinctly different environmental conditions that are unfavourable for species adapted to sky islands (Heald 1951; Smith and Farrell 2005; Robin et al. 2015; Rahim et al. 2021). Studies have highlighted the heightened sensitivity of sky island adapted taxa to extinction (e.g.Yanahan and Moore 2019; Atkins et al. 2020; Peyre et al. 2020), while also demonstrating how adaptation to sky islands may promote strong allopatric divergences (Atkins et al. 2020; Deng et al. 2020). The Western Ghats mountains in Southern India have unique sky island formations comprising high altitude laterite rich flat-topped rocky plateaus (Watve 2013). These rock outcrops have extreme environmental conditions including high temperatures during summers, heavy precipitation rains during monsoons, high wind velocity, high diurnal temperature variation, high evapotranspiration and impermeable soils (Watve 2013; Kulkarni et al. 2022). The surrounding valley and slopes tend to have less extreme abiotic environments.

We have previously investigated floral variation in two plant species in the genus *Impatiens* in the NWG rock outcrops (Rahim et al. 2021). *Impatiens lawii* is a habitat specialist species that occurs only on rock outcrops, whereas *I. oppositifolia* is a habitat generalist which co-occurs with *I. lawii* in all plateaus where the latter is found, but is found additionally in the intervening valleys. In this previous study, we focussed on three geographically proximate plateaus, and found that floral traits have diverged strongly between the three populations in *I. lawii*. In contrast, floral traits have not diverged in *I. oppositifolia*. We designed common garden experiments to test if the floral trait variation in *I. lawii* is explained by phenotypic plasticity or genetic differentiation. In these experiments, the three populations exhibited differences in floral traits that mirrored what was observed in natural populations, which suggests that the populations are genetically differentiated. There was no differentiation across populations in floral traits in *I. oppositifolia* in common garden conditions. Therefore, our results so far suggest that specialization to rock outcrops has resulted in greater inter-population divergence in *I. lawii*. However, the mechanism of floral trait differentiation is unclear. The three plateaus are geographically close to each other (separated by a maximum of 17 km), lie at approximately the same elevation (ranging from 1145 m to 1230 m), and have similar climatic conditions. Thus, it appears unlikely that local selection by abiotic factors has resulted in between-plateau floral trait divergences. Furthermore, all three plateaus have similar floral visitor communities, suggesting that diversifying selection by pollinators does not explain the floral trait variation (Rahim et al. 2021). Rahim et al (2021) concluded that genetic drift may explain the observed divergences.

The geographic proximity between populations and high dispersal rates lead to increased gene flow which, in turn, results in a weak population genetic structure because of the homogenization of allele frequencies across populations. On the other hand, when populations are geographically distant from each other and dispersal is limited, differentiation due to genetic drift is faster than the homogenizing effect of gene flow. This leads to strong structuring of gene pools and morphological traits, with a positive relationship between the degree of differentiation and spatial distance between populations – the isolation-by-distance model (Wright 1943). Thus, the isolation-by-distance model may explain the floral divergences seen in *I. lawii*.

While genetic drift can lead to similar spatial patterns of differentiation in floral and vegetative traits, selection is expected to result in different patterns of differentiation between the two types of traits. For animal pollinated plants, pollinators represent a strong selective force that shape the evolution of floral morphology (Robertson and Wyatt 1990; Johnson and Steiner 1997; Galen 1999). In contrast, abiotic factors are more important in driving variation in vegetative traits (Rico-Gray and Palacios-Rios 1996; Sapir et al. 2002; Zúñiga-Feest et al. 2015; Lambrecht et al. 2005). Examining variation in vegetative traits and the relationship between variation in floral and vegetative traits can offer important insights into the relative importance of abiotic, biotic and neutral factors that give rise to phenotypic divergence (Chalcoff et al. 2008). In our previous study, we focused only on the floral traits in the two *Impatiens* species. If the patterns of floral trait variation across populations parallel that of vegetative traits, this may be explained by neutral processes such as genetic drift or isolation-by-distance. If patterns are not congruent, it would imply that floral and vegetative traits are under differential selection regimes.

In this study, we quantified both floral and vegetative morphology across 7 plateaus representing the complete distribution of *I. lawii*. We ask the following questions

1. Do *I. lawii* populations exhibit high divergences in floral traits across the entire distribution of the species? If so, can this be explained by habitat specialization? As in Rahim et al (2021), we also quantified trait variation in *I. oppositifolia*. The explicit comparison with *I. oppositoflia* allows us to test the impact of habitat specialization.
2. Is the extent of divergence in floral and vegetative morphology correlated with geographic distance?
3. Do patterns of vegetative variation mirror that of floral trait variation across the distribution of the species?

## Methods

### Field survey and sampling

Rahim et al (2021) investigated floral variation in *I. lawii* and *I. opppositoflia* in three plateaus of the Kaas, Chalkewadi and Thoseghar. These plateaus are from the Satara cluster of rock outcrops and represent the northernmost limits of *I. lawii*. In the current study, we have surveyed the Northern Western Ghats rock formations extensively for potential plateaus with *I. lawii*. Based on this field survey, we have included four additional plateaus - Barki, Zenda, Borbet and Khamdadevi - that together with the previous three comprehensively cover the north-south and east-west distributional span of the species. We have characterized floral variation across these plateaus as well as vegetative trait variation across all seven plateaus. We have also similarly characterized floral and vegetative trait variation across the same set of plateaus in the generalist *I. oppositifoila*. Therefore, the current study enables us to compare floral and vegetative variation across 7 plateaus, and contrast patterns in the specialist and generalist species (Fig. 1).

**Fig 1.**
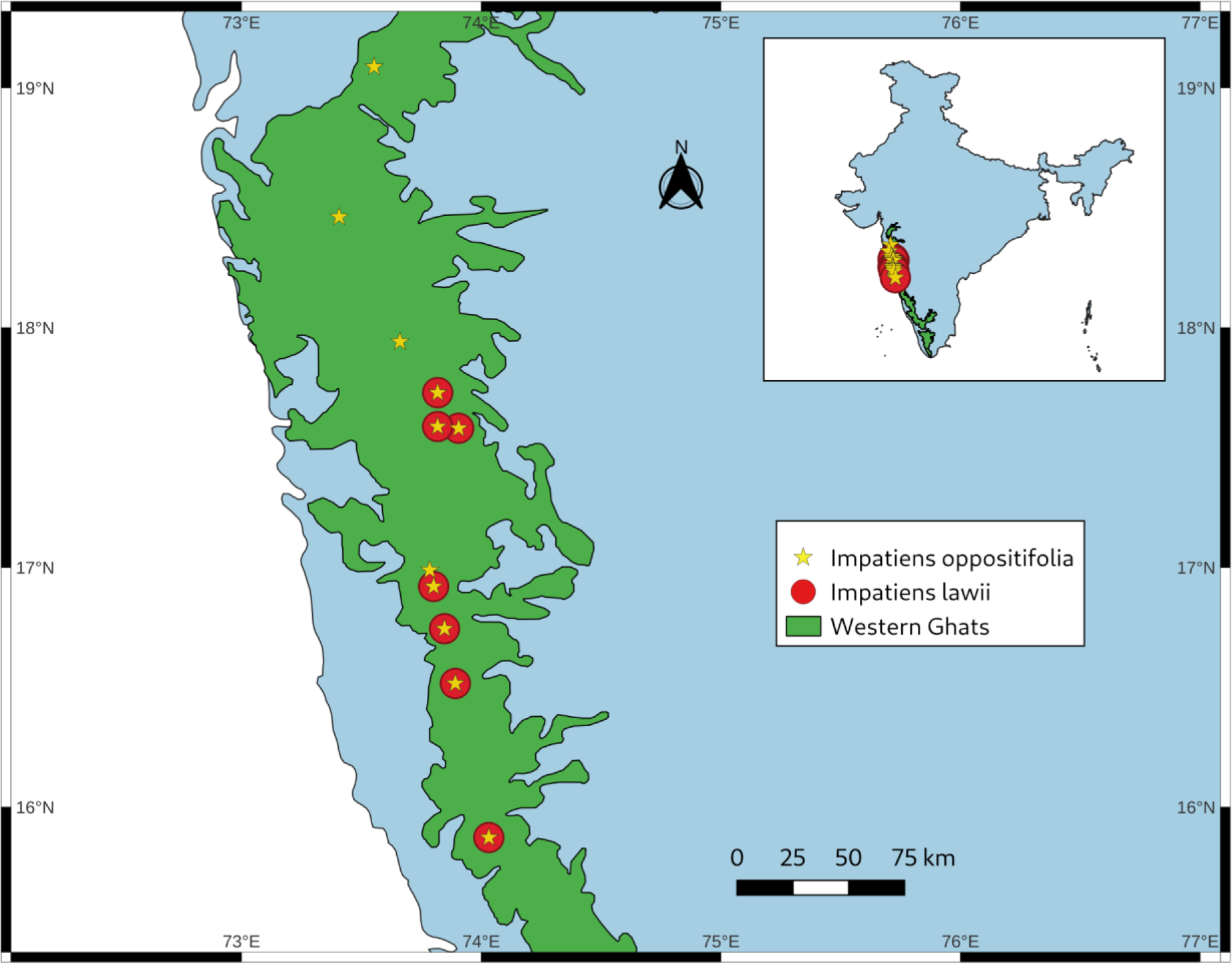
Map depicting the sampled plateau locations for specialist species *I. lawii* in northern Western Ghats. *I. oppositifolia* co-occurs with I. lawii in the seven plateaus, but is also distributed in the intervening regions.

Sampling was conducted over three years (2016-2018) during the peak flowering season (September and October). For each plateau and species, 30 individuals were collected and preserved using protocols in Rahim et al (2021). In brief, whole individuals were collected and stored in sealed plastic containers, and one flower per individual was preserved in FAA solution (Formaldehyde Alcohol Acetic Acid, 10%:50%:5% + 35% water).

### Morphometric characterization

In total, 12 floral traits and 7 vegetative traits were quantified from each individual (Table 1). All vegetative traits were measured in the field. Because flowers are small, they were dissected under a magnifying glass and floral trait variation was characterised following protocols from Rahim et al (2021) in the lab.

**Table 1.** List of morphological characters studied in populations of *Impatiens* species. All variables are measured in millimetre (mm) scale.

| Level | Code | Character |
| --- | --- | --- |
| Floral | WPL | Wing petal length |
|  | WPW | Wing petal width |
|  | STPL | Standard petal length |
|  | STPW | Standard petal width |
|  | LSPL | Lateral sepal length |
|  | LSPW | Lateral sepal width |
|  | OVL | Ovary length |
|  | OVW | Ovary width |
|  | PDL | Pedicel length |
|  | LPL | Lip length |
|  | LPW | Lip Width |
|  | SPRL | Spur length |
| Vegetative | Height | Height of the plant |
|  | NLV | Total number of leaves |
|  | NBR | Total number of branches |
|  | NFW | Total number of mature flowers |
|  | NFR | Total number of fruits |
|  | LL | Length of the leaf at its longest part |
|  | LW | Width of the leaf at its broadest part |

### Statistical analyses

All analyses were performed in R v3.5.1 (R Core Team, 2018). We used the function *prcomp* to perform Principal Component Analyses (PCA) on the respective correlation matrices for all the 22 scaled variables (Wang et al. 2016). We included all individuals in order to recognize strongly correlated factors and potential outliers (Cruz-Lustre et al. 2020). Subsequently, we excluded vegetative traits and carried out a PCA again using the 15 floral traits to explore which floral variables can best discriminate populations. PCA results were extracted and interpreted using the FactoMineR v2.4 Husson et al. 2016) and factoextra v1.0.3 (Kassambara and Mundt 2017) tools. We ran a Generalised Linear Model using the *glm* function on the first three principal components combined (PC1, PC2 and PC3), where they together constituted the dependent variable with plateau as the categorical independent variable. To test if specific plateaus differed from each other in morphological traits, we conducted pairwise comparisons through Tukey’s HSD post hoc tests with glht link function in the package multcomp v. 1.4-8 (Horthorn et al. 2008). We also fit Generalised Linear Models for each variable, including vegetative traits, separately for all populations of *I. lawii* and *I.oppositifolia*, followed by multiple comparisons of means using Tukey contrasts. The normality of residuals was checked using the Shapiro-wilk tests. To study the effect of geographical distance on trait variation, we conducted Mantel tests (Mantel, 1967) between two matrices - one comprising geographic distances between populations including all seven populations, and another comprising mean values for all 12 floral traits. The geographic distance between all population pairs was measured based on GPS coordinates of the sampled sites with the haversine formula (Nobarinezhad et al. 2019; Sinnott, 1984) using the package geosphere. The function *dist* was employed to convert the above two matrices to Euclidean distance matrices. Mantel tests were performed on these distance matrices using the Mantel test function included in the vegan package (Oksanen et al. 2015).

## Results

### Multivariate analysis

Both *I. lawii* and *I. oppositifolia* displayed moderate to strong positive correlations between most floral traits. The majority of vegetative traits were correlated with one another but not with floral traits (Appendix S1).

### Floral traits - I. lawii

In the Principal Component Analysis (PCA) of 12 floral traits in 210 *I. lawii* individuals from the seven plateaus, Principal Component (PC) PC1 explained 63.3% of the total variation with an eigenvalue of 7.6. (Appendix S2). The wing petal variables (WPL, WPW) as well as the standard petal length (STPL), keel petal variables (LPL, and LPW) were the main contributors to the PC1 and had significant positive correlations with the axis. PC2 and PC3, with corresponding eigenvalues of 1 and 0.8, explained 9% and 6.7% of the total variation, respectively. Standard petal width (STPW), spur length (SPRL) and Ovary dimensions (OVL and OVW) contributed largely to PC2. Lateral sepal width (LSPW) and size of the flower stalk or pedicel (PDL), associated with protection and optimal positioning of flowers, contributed to PC3. (Fig. 2A, Appendix S2).

**Fig 2.**
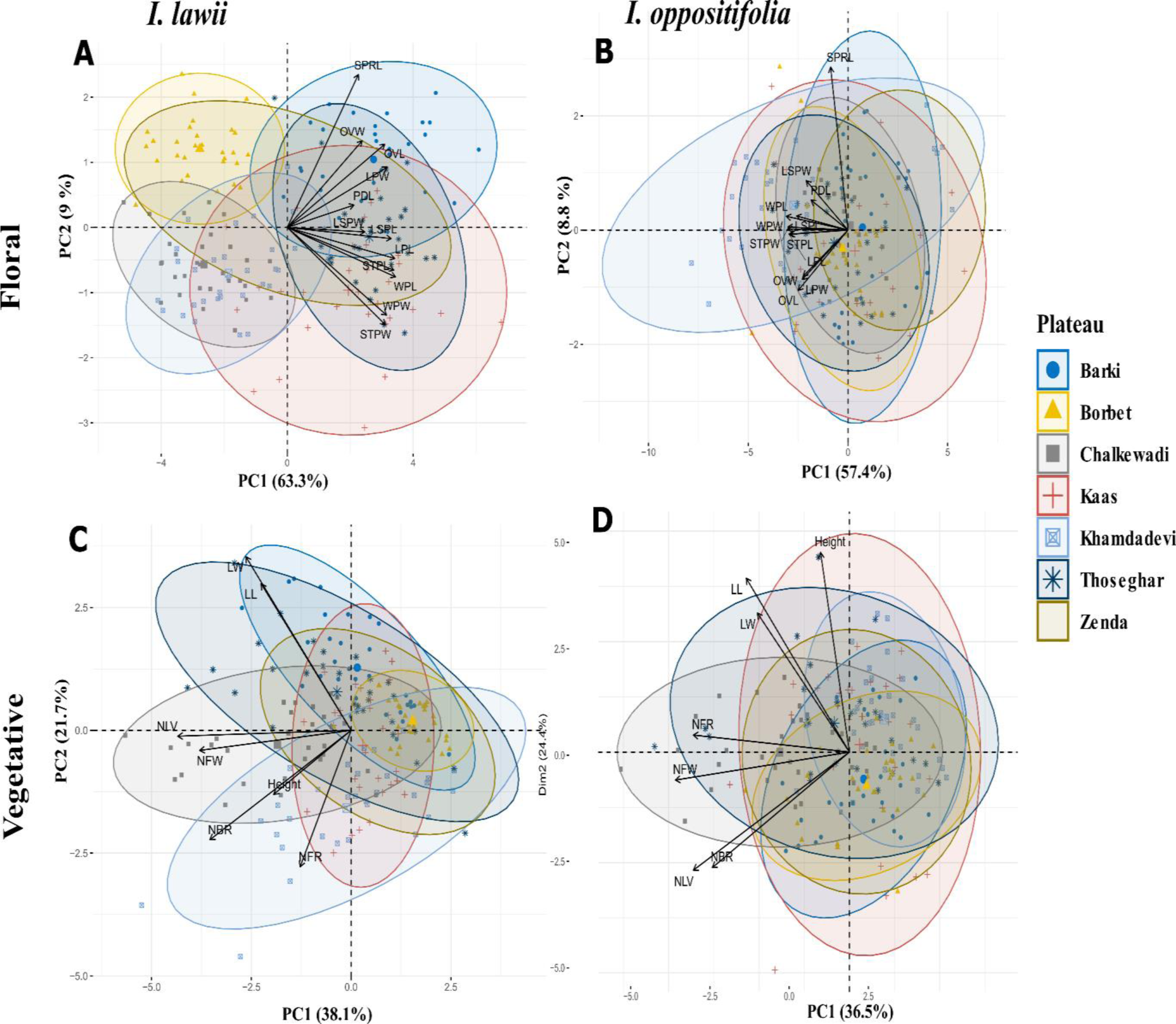
Biplots of individuals from principal component analysis (PCA) performed on 12 floral traits in *Impatiens lawii* (A) and *Impatiens oppositifolia (*B) and on 7 vegetative traits in *Impatiens lawii* (C) and *Impatiens oppositifolia (*D) for individuals from the seven plateaus. PCA Biplots are presented for 30 individuals from each species and plateau. Factor loads for PC1 and PC2 displayed as labels of the x and y-axis respectively. Ellipses indicate 95% confidence intervals. See Table 1 for character abbreviations.

The first three principal components - PC1, PC2, and PC3 - together explained 80% of the variation. Of the 21 population pairs, 18 differed significantly in floral traits in *I. lawii*, based on the GLMs and Tukey HSD post hoc tests on these three PCs jointly (Table 2, Appendix S3). Since the plateaus are distributed along a north-south latitudinal gradient (Fig. 1), we also present results of 6 comparisons of adjacent plateaus along the gradient. With the exception of the comparison between the two southernmost populations (Borbet and Khamdadevi), all populations were significantly distinct from adjacent ones (Table 3).

**Table 2.** Results of pairwise population comparisons of floral and vegetative traits in the specialist *I. lawii* and the generalist *I. oppositifolia*. A triple asterisk (***) indicates that the two plateaus were highly significantly differentiated (P < 0.001) based on a Tukey’s HSD post hoc test. Double and single asterices indicate P < 0.01 and P <0.05. Colours indicate the following: Green: P = 0.000, Blue: P < 0.001, Yellow: P < 0.01, Pink: P < 0.05, Grey: Non- significant.

|  |  | Floral |  | Vegetative |  |
| --- | --- | --- | --- | --- | --- |
|  | Plateau pair | *I. lawii* | *I. oppositifolia* | *I. lawii* | *I. oppositifolia* |
| 1 | Borbet - Barki | *** | NS | . | NS |
| 2 | Chalkewadi - Barki | *** | NS | *** | NS |
| 3 | Kaas - Barki | *** | NS | *** | NS |
| 4 | Khamdadevi - Barki | *** | *** | *** | . |
| 5 | Thoseghar - Barki | *** | NS | NS | . |
| 6 | Zenda - Barki | *** | ** | NS | NS |
| 7 | Chalkewadi - Borbet | ** | NS | *** | NS |
| 8 | Kaas - Borbet | *** | NS | *** | NS |
| 9 | Khamdadevi - Borbet | NS | NS | *** | * |
| 10 | Thoseghar - Borbet | *** | NS | ** | * |
| 11 | Zenda - Borbet | *** | *** | NS | NS |
| 12 | Kaas - Chalkewadi | *** | NS | NS | ** |
| 13 | Khamdadevi - Chalkewadi | NS | *** | ** | *** |
| 14 | Thoseghar - Chalkewadi | *** | NS | ** | *** |
| 15 | Zenda - Chalkewadi | *** | *** | ** | NS |
| 16 | Khamdadevi - Kaas | *** | *** | ** | NS |
| 17 | Thoseghar - Kaas | *** | NS | ** | NS |
| 18 | Zenda - Kaas | NS | *** | ** | NS |
| 19 | Thoseghar - Khamdadevi | *** | NS | *** | NS |
| 20 | Zenda - Khamdadevi | *** | *** | *** | * |
| 21 | Zenda - Thoseghar | . | *** | NS | * |

**Table 3.** Results of pairwise population comparisons of floral and vegetative traits in the specialist *I. lawii* and the generalist *I. oppositifolia* in comparison of of adjacent plateau pairs along the latitudinal gradient. A triple asterisk (***) indicates that the two plateaus were highly significantly differentiated (P < 0.001) based on a Tukey’s HSD post hoc test. Double and single asterices indicate P < 0.01 and P <0.05. Colours indicate the following: Green: P = 0.000, Blue: P < 0.001, Yellow: P < 0.01, Pink: P < 0.05, Grey: Non-significant.

|  | Floral |  | Vegetative |  | Floral |
| --- | --- | --- | --- | --- | --- |
|  | *I. lawii* | *I. oppositifolia* | *I. lawii* | *I. oppositifolia* | *I. lawii* |
| 1 | Kaas- Thoseghar | *** | NS | ** | NS |
| 2 | Thoseghar - Chalkewadi | *** | NS | ** | *** |
| 3 | Chalkewadi -Zenda | *** | *** | ** | NS |
| 4 | Zenda - Barki | *** | ** | NS | NS |
| 5 | Barki -Borbet | *** | NS | . | NS |
| 6 | Borbet -Khamdadevi | NS | NS | *** | * |

### *Floral traits -* I. oppositifolia

In the PCA for 12 floral traits in 210 *I. oppositifolia* individuals, PC1 explained 57.39% of the total variation with an eigenvalue of 6.88 (Appendix S2). Wing petal size (WPL, WPW), associated with floral size and pollinator attraction, as well as the variables related to pollinator efficiency - standard petal size (STPL, STPW) and keel petal length or lip length (LPL) - were the main contributors to PC1 and had significant negative correlations with the axis. PC2 and PC3, with corresponding eigenvalues of 1.06 and 0.91, explained 8.83% and 7.65% of the total variation, respectively. Length of the nectar spur (SPRL) primarily contributed to PC2 while pedicel length (PDL) contributed to PC3. Both traits were positively correlated with both axes (Fig 2B, Appendix S2).

GLM followed by a Tukey HSD based pairwise comparison of plateau based on PC1+PC2+PC3, which together accounted for 74% of total variance, showed that *I. oppositifolia* from the Zenda differed significantly from all other plateaus (Table 2, Appendix S4). Khamdadevi, the southernmost plateau in the study, differed significantly from Kaas, Chalkewadi, Zenda and Barki. The remaining 12 plateau pairs did not differ significantly from each other (Table 2, Appendix S4). In the comparison of of adjacent plateau pairs along the latitudinal gradient, Zenda differed significantly from Barki and Chalkewadi, but other pairs did not differ (Table 3).

### *Vegetative traits* - I. lawii

In the PCA of 7 vegetative traits in *I. lawii* PC1 accounted for 38.1% of the total variation with an eigenvalue of 2.66. (Appendix S5). Individual-level traits such as the number of leaves (NLV), number of branches (NBR) and number of flowers (NFW) were the main contributors to the PC1, and had significant negative correlations with the axis. PC2 and PC3, with corresponding eigenvalues of 1.5 and 1.1, explained 21.6% and 16.7% of the total variation, respectively. Leaf dimensions, such as length (LL) and width (LW), primarily contributed to PC2. Plant height was the main contributor to PC3. (Fig 2C, Appendix S5).

The first three main axes (PC1+PC2+PC3), which together accounted for 77% of the total variance, were used in the GLM, followed by pairwise analyses. These analyses revealed that in *I. lawii*, 16 of the 21 population pairs differed significantly in vegetative traits (Table 2, Appendix S6). Comparisons of adjacent population pairs along the latitudinal gradient showed that all pairs except Barki-Zenda are significantly different (Table 3).

### *Vegetative traits* - I. oppositifolia

In the PCA of 7 vegetative traits in *I. oppositifolia*, PC1 accounted for 36.46% of the total variance with an eigenvalue of 2.5. (Appendix S5). Individual-level traits such as the number of flowers (NFW), number of fruits (NFR), number of leaves (NLV), and number of branches (NBR) were the main contributors to the PC1. They had significant negative correlations with the axis. PC2 and PC3, with corresponding eigenvalues of 1.71 and 0.82, explained 24.43% and 11.84% of the total variation, respectively. Plant height and leaf length (LL) primarily contributed to PC2 while the height of the plant along with leaf width dimension (LW) contributed to PC3. Both traits were positively correlated with both axes (Fig 2D, Appendix S5).

PC1, PC2 and PC3 together accounted for 73% of the variation. GLM followed by a pairwise post-hoc analyses showed that 9 of the 21 population pairs were differed significantly from each other (Table 2, Appendix S7). Of the 6 north-south adjacent pairs, only the comparison between Thoseghar-Chalkewadi and that between Borbet-Khamdadevi was significant (Table 3).

### Correlation of trait variation with geographic distance and altitude

Mantel tests based on Pearson’s product-moment correlation revealed that the distance between the plateaus’ geographical locations did not correlate with differences in both floral traits (Mantel statistic r= -0.0507, p= 0.46) (Fig 3A) and vegetative traits (Mantel statistic r= -0.2166, p= 0.78) (Fig 3C) in *I. lawii.* Similarly, altitudinal differences between plateaus were also not correlated with differences in floral (Mantel statistic r= -0.079, p= 0.57) (Fig 4A) or vegetative (Mantel statistic r= -0245, p= 0.81) traits (Fig 4C) in *I. lawii* (Table 4).

**Fig 3.**
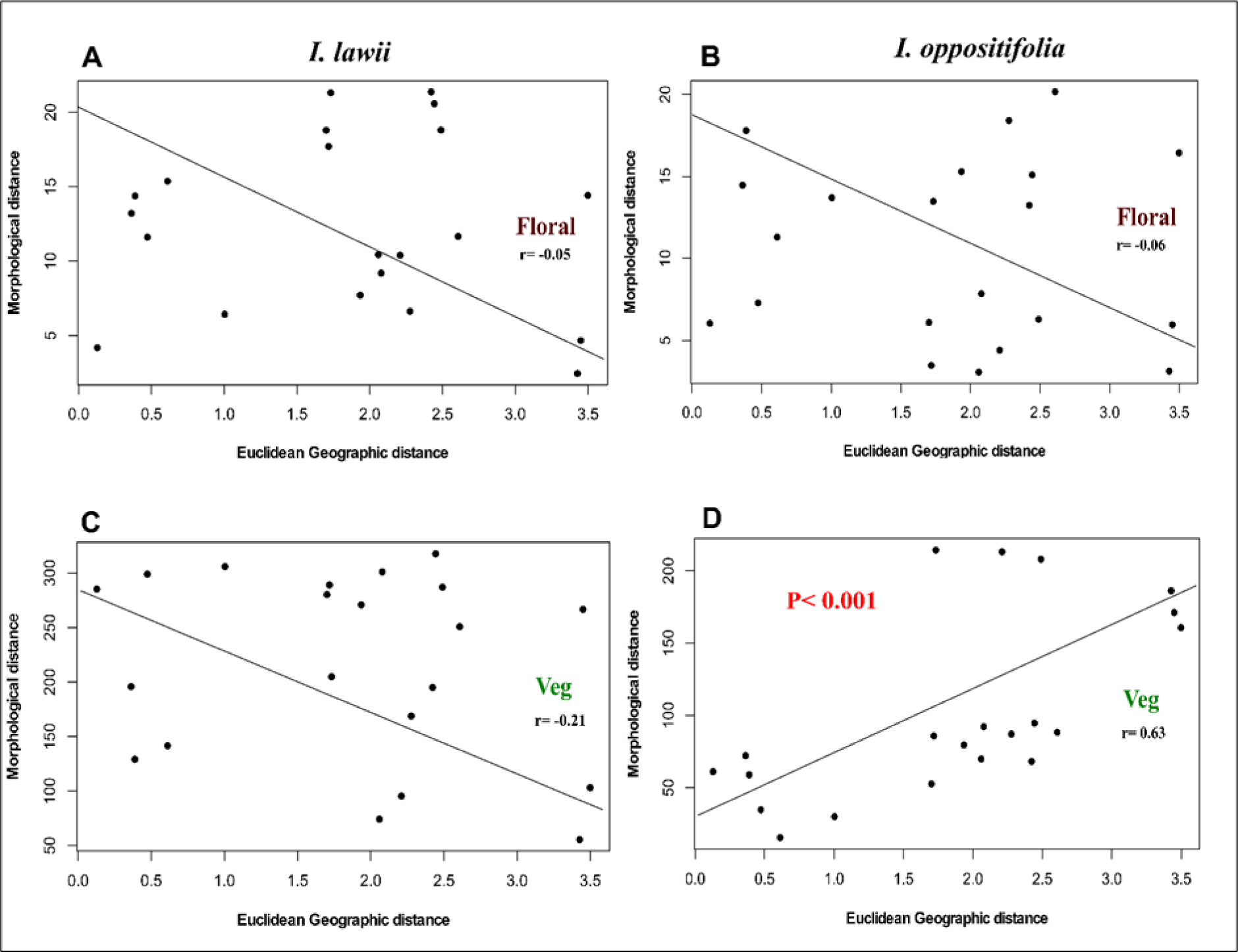
Results of Mantel tests for correlations of morphological distance with geographic distance. Correlation of floral traits with geographic distance in A) *I. lawii*; B*) I. oppositifolia*Correlation of vegetative traits with geographic distance in C) *I. lawii*; D*) I. oppositifolia*.

**Fig 4.**
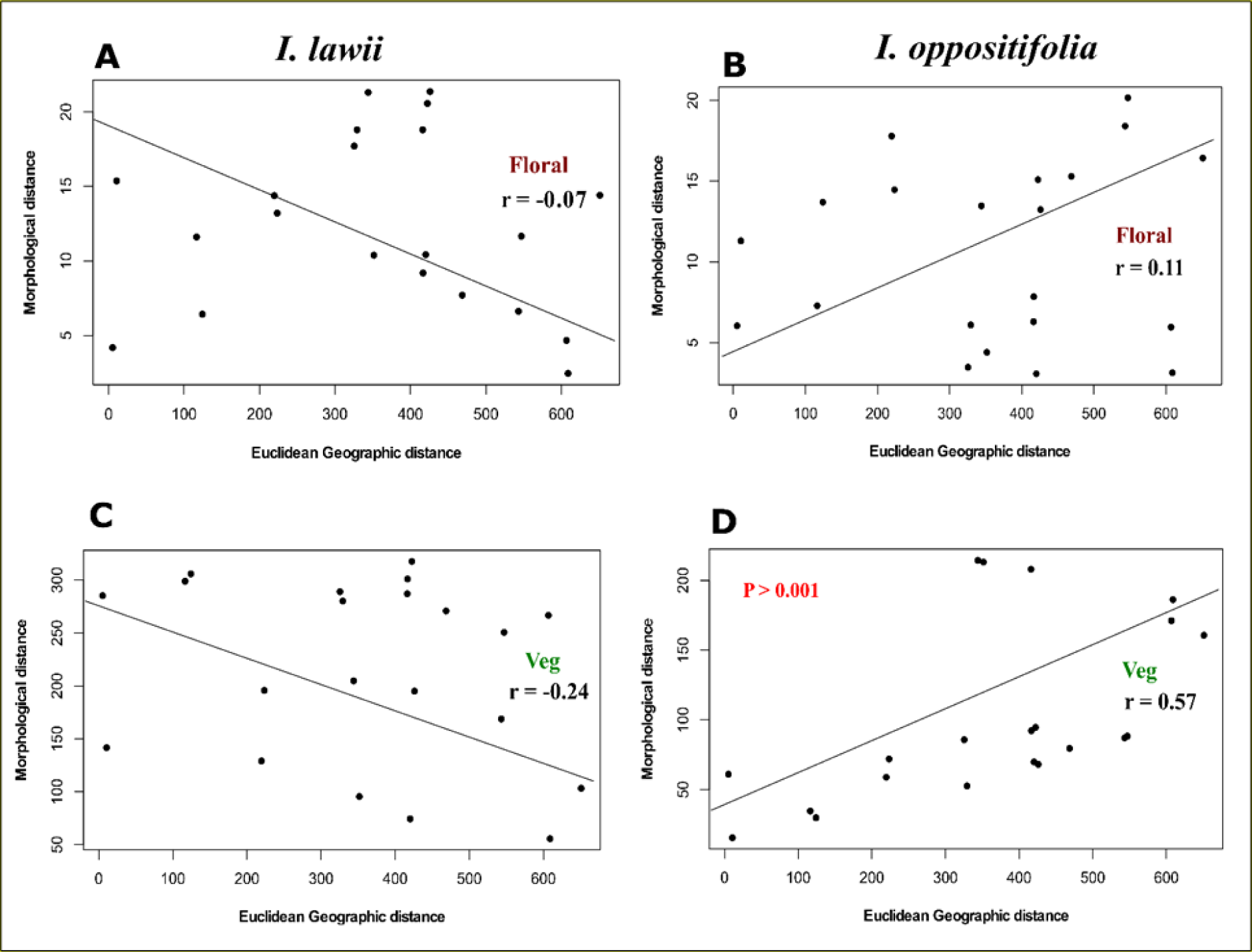
Results of Mantel tests for correlations of morphological distance with altitudinal distance. Correlation of floral traits with altitude distance in A) *I. lawii*; B*) I. oppositifolia.* Correlation of vegetative traits with altitude distance in C) *I. lawii*; D*) I. oppositifolia*.

**Table 4.** Results of Mantel tests for correlations of morphological distance with geographic distance and altitude. Values reported are Mantel statistic r, which ranges from - 1 to 1. Values close to -1 indicate a strong negative correlation, values close to 1 indicate strong positive correlation, and; values close to 0 indicate no correlation. Comparisons with significant correlations (P<0.001) are shown in green.

|  | Morphological distance v/s<br>Geographical distance |  | Morphological distance v/s<br>Altitude distance |  |
| --- | --- | --- | --- | --- |
|  | *I. lawii* | *I. oppositifolia* | *I. lawii* | *I. oppositifolia* |
| Floral | -0.05 | -0.06 | -0.07 | 0.11 |
| Veg | -0.21 | 0.63 | -0.24 | 0.57 |

However, in *I. oppositifolia*, the distance between plateaus was significantly correlated with differences in vegetative traits (Mantel statistic r= 0.639, p= 0.01) (Fig 3D) but not floral traits (Mantel statistic r= -0.066, p= 0.44) (Fig 3B). Altitudinal differences between were significantly correlated with differences in vegetative (Mantel statistic r= 0.572, p= 0.01) traits (Fig 4D), but not floral traits (Mantel statistic r= 0.110, p= 0.35) (Fig 4B).

## Discussion

Floral traits show consistently deeper divergences between population pairs in *I. lawii* than in *I. oppositfolia*. Visual inspection of the floral PCA plots (Fig 2) indicate that the plateaus group into respective clusters in *I. lawii*, while there is considerably stronger overlap among plateaus in the case of *I. oppositifolia*. Statistical comparisons based on the first three PCs of floral traits reveal that ca. 85% (17 of 21) of pairwise plateau comparisons were significant (Table 2). On the other hand, only about 42% of plateau pairs (9 of 21) were significantly differentiated in *I. lawii* (Table 2). Furthermore, when comparing adjacent plateaus along the latitudinal gradient, 5 of 6 plateau pairs in *I. lawii* were significantly different, but only 3 pairs in *I. oppositifolia* were (Table 2). Therefore, we conclude that specialization to rock outcrop habitats has led to deeper allopatric divergences in floral morphology across the distribution of *I. lawii*.

The among population variation in morphological traits can be due to (i) phenotypic plasticity (ii) genetic differentiation, or (iii) a combination of (i) and (ii). Results from the common garden experiments in our previous study (Rahim et al. 2021) suggest that phenotypic plasticity does not explain floral trait variation within the three plateaus of the Satara cluster. These plateaus are at similar altitudes, and geographically proximate, with the maximum distance between two plateaus not more than 17km (Rahim et al. 2021). Our current study includes a larger geographic sampling, with > 200 km distance between plateaus (Appendix S8-S9) and there is variation in altitude across the plateaus. Therefore, we cannot rule out the influence of plasticity. We surmise that genetic differentiation *per se* or in combination with plasticity explains cross-population floral trait variation in the two *Impatiens* species observed in our study.

### Floral versus vegetative traits

In *I. lawii*, vegetative traits were less strongly differentiated across plateaus than were floral traits (Table 2). Moreover, there was no consistency in the pattern of structuring of variation across plateaus. For instance, Khamdadevi - Chalkewadi, Zenda-Kaas and Zenda-Thoseghar were not significantly differentiated in floral traits. Surprisingly, these population pairs were among the pairs that were highly significantly differentiated in vegetative traits. Therefore, we conclude that the processes leading to differentiation across populations in this species varies between vegetative and floral traits. Such decoupling of floral and vegetative trait variation has been reported widely in many plants (Berg 1959; Pélabon et al. 2011; Conner and Lande 2014).

Under an isolation-by-distance model (Wright 1943), genetic drift drives population genetic structuring. This model predicts an association between geographic distance and phenotypic distance. The Mantel tests indicate no such association for floral traits in both *I. lawii* and *I. oppositifolia* (Fig 4; Table 4), suggesting that genetic drift alone cannot explain floral trait variation in these species. Our previous work on the three plateaus (Rahim et al. 2021) suggests that two *Impatiens* species are not specialized in their pollination niche. Although there were differences in floral visitation rates between plateaus, both species are pollinated by a large number of pollinator groups. Thus, it is unlikely that geographically patterned selection by different pollinators explains floral trait variation across the range of these two species.

Interestingly, there was a significant correlation between geographic distance and vegetative trait variation in *I. oppositifolia*, but not in *I. lawii* (Fig 4D; Table 2). This result by itself suggests a role of genetic drift in *I. oppositifolia*. However, genetic drift should result in consistency in patterns of variation of floral and vegetative traits. There was no consistency in these patterns - plateaus that differed in floral traits did not necessarily differ in vegetative traits (Table 2). Furthermore, vegetative traits in *I. oppositifolia* are also correlated with altitude, suggesting selection by the abiotic factors that vary with altitude. Studies on several plants have indicated that geographic variation in vegetative morphology is driven by variation in climatic (Joshi et al. 2001; Santamaría et al. 2003; Macel et al. 2007; Baughman et al. 2019) and edaphic parameters (Antonovics et al. 1971; Schat et al. 1996; Wright et al. 2006). Factors such as aridity and temperature, which tend to be associated with elevation, may act differentially on vegetative morphology across plateaus in *I. oppositifolia*, leading to local adaptation. If abiotic factors select for similar traits across both species, differences in the abiotic environment between plateaus are expected to lead to a stronger phenological structuring in the specialist species (compared to the generalist) via local adaptation. This is because geneflow between populations in the generalist will override the effect of local adaptation. Therefore, the absence of an association between vegetative variation and altitude in the specialist species is intriguing.

Although floral morphology in animal pollinated plants has often been presumed to be primarily selected by pollinators, many studies have highlighted the importance of selection by abiotic factors (Strauss and Whittall 2006; Arista et al. 2013; Dalrymple et al. 2020). Selection on vegetative traits can indirectly affect floral traits (Galen 1999; Lambrecht and Dawson 2007). Because there is no congruence between the pattern of floral and vegetative variation across the range of *I. lawii*, an indirect influence on floral traits by direct selection on vegetative traits seems unlikely. Multiple abiotic factors, including edaphic factors, are known to directly select for particular floral traits (Chalker-Scott 1999; Arista et al. 2013; Koski and Ashman 2015; Keefover-Ring et al. 2022; reviewed in Caruso et al. 2019; Dalrymple et al. 2020). The northern Western Ghats rocky plateaus originated ca. 65 million through Deccan Trap volcanism, followed by chemical leaching of the parent rock in high rainfall conditions and hardening that eventually led to the formation of laterite (Watve 2008). The soil is shallow, sandy to sandy loam in texture, phosphate poor and acidic (Watve 2008). The soils have low nutrient availability (Kulkarni et al. 2022). Little information exists about variation in edaphic properties across plateaus in the current study. We speculate that variation in soil and rock properties may explain some of the floral trait variation in *I. lawii*.

### Summary and Conclusions

Our results indicate that floral trait variation in both the rocky outcrop habitat specialist *I. lawii* and the generalist *I. oppositifolia* are geographically structured. Floral trait variation was more strongly structured in the specialist, with more population pairs differing significantly from each other. These results support the idea that habitat specialization has led to stronger between population divergences in the specialist. In both species, vegetative traits were more strongly structured across plateaus than were floral traits. This may be because of location adaptation due to selection by abiotic factors. In the generalist, differences in vegetative traits between plateaus are correlated with geograhic and elevational differences, further supporting the role of selection by abiotic factors. It will be interesting to analyze the population genetic structure of the two species, and explore how genetic differentiation relates to floral, vegetative, and environmental differences between plateaus.

## Acknowledgements

We thank Dr. Deepak Barua for logistic support, discussions and for assistance in error checking data set. We also acknowledge the assistance of Dr. Gopal Murali in the GLM analysis, as well as the fieldwork assistance of Ravi Umadi and Sairandhri Lapalikar in gathering morphological data

## Appendices

**Appendix S1.** Correlation plot showing correlation between vegetative and floral traits in *I. lawii* and between floral traits in *I. lawii* and *I.oppositifolia.* All floral traits are moderate – high, positively correlated to each other in both species. Vegetative traits are not correlated to floral traits. See Table 1 for character abbreviations.

**Appendix S2.** Correlation between floral variables and first three principal component axes with eigenvalues (EV), variance percent (Var) and cumulative variance percent (CVA) derived from Principal Component Analysis (PCA) of the seven populations of *I. lawii* and *I. oppositifolia* in wild. See Table 1 for character abbreviations.

**Appendix S3.** Results of GLM on first three PC’s (PC1+PC2+PC3) followed by Tukey HSD posthoc test in *I. lawii* conducted for 12 floral traits. A triple asterisk (***) indicates that the two plateaus were highly significantly differentiated (P < 0.001) based on a Tukey’s HSD post hoc test. Double and single asterices indicate P < 0.01 and P <0.05. Colours indicate the following: Green: P = 0.000, Blue: P < 0.001, Yellow: P < 0.01, Pink: P < 0.05, Grey: Non-significant.

**Appendix S4.** Results of GLM on first three PC’s (PC1+PC2+PC3) followed by Tukey HSD posthoc test in *I. oppositifolia* conducted for 12 floral traits. A triple asterisk (***) indicates that the two plateaus were highly significantly differentiated (P < 0.001) based on a Tukey’s HSD post hoc test. Double and single asterices indicate P < 0.01 and P <0.05. Colours indicate the following: Green: P = 0.000, Blue: P < 0.001, Yellow: P < 0.01, Pink: P < 0.05, Grey: Non-significant.

**Appendix S5.** Correlation between vegetative variables and first three principal component axes with eigenvalues (EV), variance percent (Var) and cumulative variance percent (CVA) derived from Principal Component Analysis (PCA) of the seven populations of *I. lawii* and *I. oppositifolia* in wild. See Table 1 for character abbreviations.

**Appendix S6.** Results of GLM on first three PC’s (PC1+PC2+PC3) followed by Tukey HSD posthoc test in *I. lawii* conducted for 7 vegetative traits. A triple asterisk (***) indicates that the two plateaus were highly significantly differentiated (P < 0.001) based on a Tukey’s HSD post hoc test. Double and single asterices indicate P < 0.01 and P <0.05. Colours indicate the following: Green: P = 0.000, Blue: P < 0.001, Yellow: P < 0.01, Pink: P < 0.05, Grey: Non-significant

**Appendix S7.** Results of GLM on first three PC’s (PC1+PC2+PC3) followed by Tukey HSD posthoc test in *I. oppositifolia* conducted for 7 vegetative traits. A triple asterisk (***) indicates that the two plateaus were highly significantly differentiated (P < 0.001) based on a Tukey’s HSD post hoc test. Double and single asterices indicate P < 0.01 and P <0.05. Colours indicate the following: Green: P = 0.000, Blue: P < 0.001, Yellow: P < 0.01, Pink: P < 0.05, Grey: Non-significant.

**Appendix S8.** Table representing the distance between Plateaus (Km) in the present study. The minimum and maximum distance between two plateaus are approximately 7 Km and 200 Km respectively.

**Appendix S9.** Table representing the details of the study area.

