## Supplementary Data for "Drivers of phenotypic differentiation across rock outcrop sky islands in *Impatiens* plants"

**Appendices**

**Appendix S1**: Correlation plot showing correlation between vegetative and floral traits in *I. lawii* and between floral traits in *I. lawii* and *I.oppositifolia.* All floral traits are moderate –high, positively correlated to each other in both species. Vegetative traits are not correlated to floral traits. See Table 1 for character abbreviations.

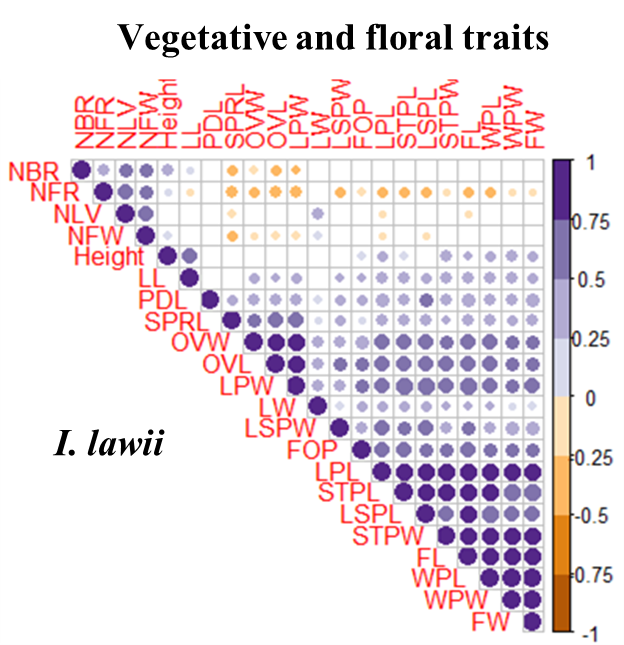

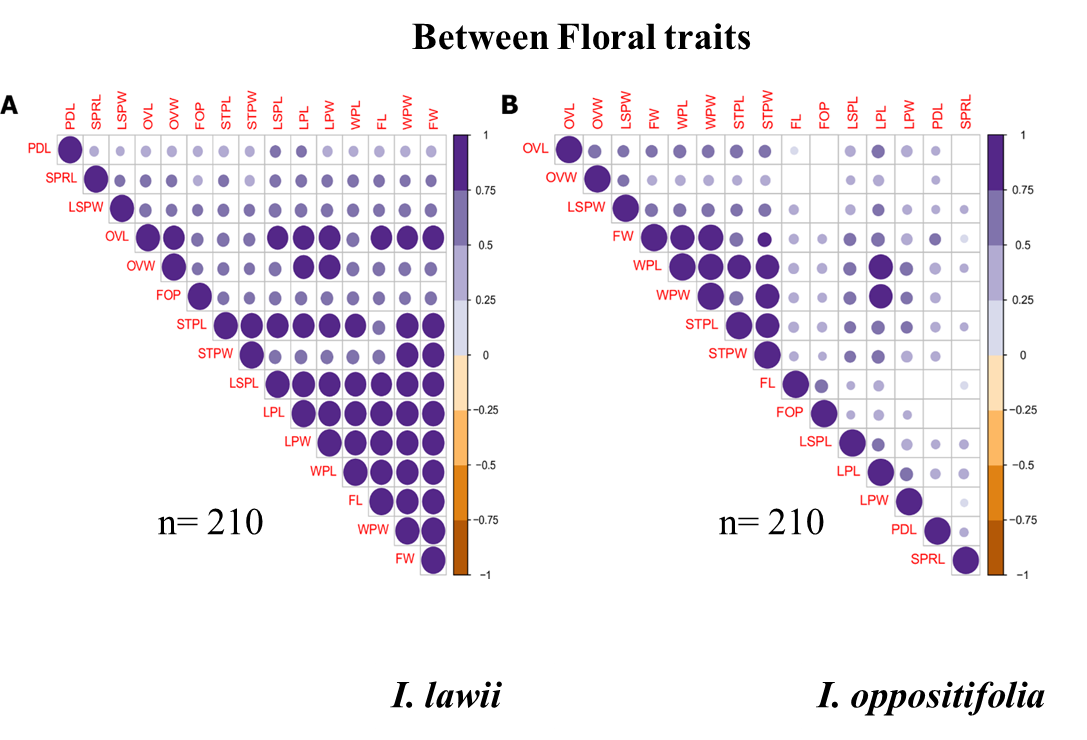

**Appendix S2:** Correlation between floral variables and first three principal component axes with eigenvalues (EV), variance percent (Var) and cumulative variance percent (CVA) derived from Principal Component Analysis (PCA) of the seven populations of *I. lawii* and *I. oppositifolia* in wild. See Table 1 for character abbreviations.

| **Species** | **Trait** | **PC** | | |
| --- | --- | --- | --- | --- |
|  |  | **PC1** | **PC2** | **PC3** |
| **a) *I. lawii*** | WPL | 0.92 | -0.20 | 0.09 |
|  | WPW | 0.84 | -0.36 | 0.07 |
|  | STPL | 0.90 | -0.18 | -0.04 |
|  | STPW | 0.83 | -0.40 | 0.04 |
|  | LSPL | 0.88 | -0.05 | -0.11 |
|  | LSPW | 0.66 | -0.02 | -0.43 |
|  | OVL | 0.83 | 0.34 | 0.10 |
|  | OVW | 0.64 | 0.36 | -0.48 |
|  | PDL | 0.57 | 0.09 | 0.57 |
|  | LPL | 0.91 | -0.13 | 0.00 |
|  | LPW | 0.85 | 0.25 | 0.03 |
|  | SPRL | 0.60 | 0.63 | 0.19 |
|  | EV | 7.60 | 1.09 | 0.81 |
|  | Var % | 63.33 | 9.05 | 6.77 |
|  | CV % | 63.33 | 72.38 | 79.15 |
| **b) *I. oppositofolia*** | WPL | -0.93 | 0.07 | -0.18 |
|  | WPW | -0.90 | 0.02 | -0.16 |
|  | STPL | -0.87 | -0.02 | -0.20 |
|  | STPW | -0.91 | 0.01 | -0.20 |
|  | LSPL | -0.77 | 0.07 | -0.18 |
|  | LSPW | -0.63 | 0.26 | -0.36 |
|  | OVL | -0.75 | -0.32 | 0.36 |
|  | OVW | -0.69 | -0.26 | 0.23 |
|  | PDL | -0.55 | 0.16 | 0.54 |
|  | LPL | -0.89 | -0.02 | 0.10 |
|  | LPW | -0.67 | -0.24 | 0.21 |
|  | SPRL | -0.26 | 0.85 | 0.30 |
|  | **EV** | 6.89 | 1.06 | 0.92 |
|  | **Var %** | 57.39 | 8.84 | 7.66 |
|  | **CV%** | 57.39 | 66.23 | 73.88 |

**Appendix S3:** Results of GLM on first three PC's (PC1+PC2+PC3) followed by Tukey HSD posthoc test in *I. lawii* conducted for 12 floral traits.A triple asterisk (***) indicates that the two plateaus were highly significantly differentiated (P < 0.001) based on a Tukey’s HSD post hoc test. Double and single asterices indicate P < 0.01 and P <0.05. Colours indicate the following: Green:  P = 0.000, Blue: P < 0.001, Yellow: P < 0.01, Pink: P < 0.05, Grey: Non-significant.

| **Plateau comparison** | **Estimate** | **Std. Error** | **z value** | **Pr(>\|z\|)** | **Sig** |
| --- | --- | --- | --- | --- | --- |
| Borbet - Barki | -5.9916 | 0.4319 | -13.873 | < 0.001 | ******* |
| Chalkewadi - Barki | -7.5273 | 0.4246 | -17.727 | < 0.001 | ******* |
| Kaas - Barki | -3.6895 | 0.4246 | -8.689 | < 0.001 | ******* |
| Khamdadevi - Barki | -6.3971 | 0.4281 | -14.941 | < 0.001 | ******* |
| Thoseghar - Barki | -1.7741 | 0.4358 | -4.07 | < 0.001 | ******* |
| Zenda - Barki | -3.3605 | 0.5383 | -6.243 | < 0.001 | ******* |
| Chalkewadi - Borbet | -1.5357 | 0.4319 | -3.556 | 0.00676 | ****** |
| Kaas - Borbet | 2.3021 | 0.4319 | 5.33 | < 0.001 | ******* |
| Khamdadevi - Borbet | -0.4055 | 0.4353 | -0.931 | 0.96708 | **NS** |
| Thoseghar - Borbet | 4.2175 | 0.4429 | 9.522 | < 0.001 | ******* |
| Zenda - Borbet | 2.6312 | 0.544 | 4.836 | < 0.001 | ******* |
| Kaas - Chalkewadi | 3.8379 | 0.4246 | 9.038 | < 0.001 | ******* |
| Khamdadevi - Chalkewadi | 1.1303 | 0.4281 | 2.64 | 0.11251 | **NS** |
| Thoseghar - Chalkewadi | 5.7533 | 0.4358 | 13.2 | < 0.001 | ******* |
| Zenda - Chalkewadi | 4.1669 | 0.5383 | 7.741 | < 0.001 | ******* |
| Khamdadevi - Kaas | -2.7076 | 0.4281 | -6.324 | < 0.001 | ******* |
| Thoseghar - Kaas | 1.9154 | 0.4358 | 4.395 | < 0.001 | ******* |
| Zenda - Kaas | 0.329 | 0.5383 | 0.611 | 0.99643 | **NS** |
| Thoseghar - Khamdadevi | 4.623 | 0.4393 | 10.524 | < 0.001 | ******* |
| Zenda - Khamdadevi | 3.0366 | 0.5411 | 5.612 | < 0.001 | ******* |
| Zenda - Thoseghar | -1.5864 | 0.5472 | -2.899 | 0.05649 | **.** |

**Appendix S4:** Results of GLM on first three PC's (PC1+PC2+PC3) followed by Tukey HSD posthoc test in *I. oppositifolia* conducted for 12 floral traits. A triple asterisk (***) indicates that the two plateaus were highly significantly differentiated (P < 0.001) based on a Tukey’s HSD post hoc test. Double and single asterices indicate P < 0.01 and P <0.05. Colours indicate the following: Green:  P = 0.000, Blue: P < 0.001, Yellow: P < 0.01, Pink: P < 0.05, Grey: Non-significant.

| **Plateau comparison** | **Estimate** | **Std. Error** | **z value** | **Pr(>\|z\|)** | **Sig** |
| --- | --- | --- | --- | --- | --- |
| Borbet - Barki | -1.95039 | 0.64038 | -3.046 | 0.076 | **NS** |
| Chalkewadi - Barki | -0.63492 | 0.67163 | -0.945 | 0.9651 | **NS** |
| Kaas - Barki | -0.73344 | 0.65171 | -1.125 | 0.9204 | **NS** |
| Khamdadevi - Barki | -3.4138 | 0.64587 | -5.286 | <0.001 | ******* |
| Thoseghar - Barki | -1.25456 | 0.65792 | -1.907 | 0.4753 | **NS** |
| Zenda - Barki | 2.50381 | 0.65171 | 3.842 | 0.0022 | ****** |
| Chalkewadi - Borbet | 1.31547 | 0.67163 | 1.959 | 0.441 | **NS** |
| Kaas - Borbet | 1.21696 | 0.65171 | 1.867 | 0.5019 | **NS** |
| Khamdadevi - Borbet | -1.46341 | 0.64587 | -2.266 | 0.2604 | **NS** |
| Thoseghar - Borbet | 0.69583 | 0.65792 | 1.058 | 0.9402 | **NS** |
| Zenda - Borbet | 4.4542 | 0.65171 | 6.835 | <0.001 | ******* |
| Kaas - Chalkewadi | -0.09852 | 0.68245 | -0.144 | 1 | **NS** |
| Khamdadevi - Chalkewadi | -2.77888 | 0.67688 | -4.105 | <0.001 | ******* |
| Thoseghar - Chalkewadi | -0.61964 | 0.68838 | -0.9 | 0.9726 | **NS** |
| Zenda - Chalkewadi | 3.13873 | 0.68245 | 4.599 | <0.001 | ******* |
| Khamdadevi - Kaas | -2.68037 | 0.65711 | -4.079 | <0.001 | ******* |
| Thoseghar - Kaas | -0.52112 | 0.66896 | -0.779 | 0.987 | **NS** |
| Zenda - Kaas | 3.23725 | 0.66285 | 4.884 | <0.001 | ******* |
| Thoseghar - Khamdadevi | 2.15924 | 0.66328 | 3.255 | 0.0795 | **NS** |
| Zenda - Khamdadevi | 5.91761 | 0.65711 | 9.005 | <0.001 | ******* |
| Zenda - Thoseghar | 3.75837 | 0.66896 | 5.618 | <0.001 | ******* |

**Appendix S5:** Correlation between vegetative variables and first three principal component axes with eigenvalues (EV), variance percent (Var) and cumulative variance percent (CVA) derived from Principal Component Analysis (PCA) of the seven populations of *I. lawii* and *I. oppositifolia* in wild. See Table 1 for character abbreviations.

| **Species** | **Trait** | **PC** | | |
| --- | --- | --- | --- | --- |
|  |  | **PC1** | **PC2** | **PC3** |
| **a) *I. lawii*** | LL | -0.46 | 0.61 | -0.51 |
|  | LW | -0.54 | 0.73 | -0.05 |
|  | Height | -0.40 | -0.27 | -0.65 |
|  | NBR | -0.73 | -0.46 | 0.07 |
|  | NLV | -0.89 | -0.02 | 0.23 |
|  | NFW | -0.78 | -0.08 | 0.48 |
|  | NFR | -0.26 | -0.57 | -0.45 |
|  | EV | 2.67 | 1.52 | 1.17 |
|  | Var % | 38.11 | 21.68 | 16.72 |
|  | CV % | 38.11 | 59.79 | 76.50 |
| **b) *I. oppositofolia*** | LL | -0.48 | 0.66 | -0.18 |
|  | LW | -0.43 | 0.53 | 0.45 |
|  | Height | -0.13 | 0.76 | -0.51 |
|  | NBR | -0.64 | -0.44 | -0.41 |
|  | NLV | -0.73 | -0.45 | -0.19 |
|  | NFW | -0.81 | -0.11 | 0.18 |
|  | NFR | -0.73 | 0.06 | 0.31 |
|  | **EV** | 2.55 | 1.71 | 0.83 |
|  | **Var %** | 36.46 | 24.44 | 11.85 |
|  | **CV%** | 36.46 | 60.90 | 72.75 |

**Appendix S6:** Results of GLM on first three PC's (PC1+PC2+PC3) followed by Tukey HSD posthoc test in *I. lawii* conducted for 7 vegetative traits.A triple asterisk (***) indicates that the two plateaus were highly significantly differentiated (P < 0.001) based on a Tukey’s HSD post hoc test. Double and single asterices indicate P < 0.01 and P <0.05. Colours indicate the following: Green:  P = 0.000, Blue: P < 0.001, Yellow: P < 0.01, Pink: P < 0.05, Grey: Non-significant.

| **Plateau comparison** | **Estimate** | **Std. Error** | **z value** | **Pr(>\|z\|)** | **Sig** |
| --- | --- | --- | --- | --- | --- |
| Borbet - Barki | 1.31344 | 0.44854 | 2.928 | 0.05246 | **.** |
| Chalkewadi - Barki | -1.89879 | 0.44485 | -4.268 | < 0.001 | ******* |
| Kaas - Barki | -1.91753 | 0.44485 | -4.31 | < 0.001 | ******* |
| Khamdadevi - Barki | -3.49309 | 0.44485 | -7.852 | < 0.001 | ******* |
| Thoseghar - Barki | -0.32967 | 0.45246 | -0.729 | 0.99076 | **NS** |
| Zenda - Barki | 0.20301 | 0.53913 | 0.377 | 0.99977 | **NS** |
| Chalkewadi - Borbet | -3.21223 | 0.44854 | -7.161 | < 0.001 | ******* |
| Kaas - Borbet | -3.23097 | 0.44854 | -7.203 | < 0.001 | ******* |
| Khamdadevi - Borbet | -4.80653 | 0.44854 | -10.716 | < 0.001 | ******* |
| Thoseghar - Borbet | -1.6431 | 0.45609 | -3.603 | 0.00568 | ****** |
| Zenda - Borbet | -1.11043 | 0.54218 | -2.048 | 0.38139 | **NS** |
| Kaas - Chalkewadi | -0.01874 | 0.44485 | -0.042 | 1 | **NS** |
| Khamdadevi - Chalkewadi | -1.5943 | 0.44485 | -3.584 | 0.00608 | ****** |
| Thoseghar - Chalkewadi | 1.56913 | 0.45246 | 3.468 | 0.00938 | ****** |
| Zenda - Chalkewadi | 2.1018 | 0.53913 | 3.899 | 0.00187 | ****** |
| Khamdadevi - Kaas | -1.57556 | 0.44485 | -3.542 | 0.00734 | ****** |
| Thoseghar - Kaas | 1.58787 | 0.45246 | 3.509 | 0.00796 | ****** |
| Zenda - Kaas | 2.12054 | 0.53913 | 3.933 | 0.00149 | ****** |
| Thoseghar - Khamdadevi | 3.16343 | 0.45246 | 6.992 | < 0.001 | ******* |
| Zenda - Khamdadevi | 3.6961 | 0.53913 | 6.856 | < 0.001 | ******* |
| Zenda - Thoseghar | 0.53268 | 0.54542 | 0.977 | 0.95864 | **NS** |

**Appendix S7:** Results of GLM on first three PC's (PC1+PC2+PC3) followed by Tukey HSD posthoc test in *I. oppositifolia* conducted for 7 vegetative traits.A triple asterisk (***) indicates that the two plateaus were highly significantly differentiated (P < 0.001) based on a Tukey’s HSD post hoc test. Double and single asterices indicate P < 0.01 and P <0.05. Colours indicate the following: Green:  P = 0.000, Blue: P < 0.001, Yellow: P < 0.01, Pink: P < 0.05, Grey: Non-significant.

| Plateau comparison | Estimate | Std. Error | z value | Pr(>\|z\|) | Sig |
| --- | --- | --- | --- | --- | --- |
| Borbet - Barki | -0.20199 | 0.5406 | -0.374 | 0.99979 | **NS** |
| Chalkewadi - Barki | -1.16662 | 0.5406 | -2.158 | 0.3187 | **NS** |
| Kaas - Barki | 0.93257 | 0.54516 | 1.711 | 0.60866 | **NS** |
| Khamdadevi - Barki | 1.57614 | 0.5363 | 2.939 | 0.05128 | **.** |
| Thoseghar - Barki | 1.53017 | 0.54516 | 2.807 | 0.07427 | **.** |
| Zenda - Barki | -0.17819 | 0.55001 | -0.324 | 0.99991 | **NS** |
| Chalkewadi - Borbet | -0.96463 | 0.536 | -1.8 | 0.54818 | **NS** |
| Kaas - Borbet | 1.13456 | 0.5406 | 2.099 | 0.35309 | **NS** |
| Khamdadevi - Borbet | 1.77813 | 0.53166 | 3.344 | 0.01445 | ***** |
| Thoseghar - Borbet | 1.73216 | 0.5406 | 3.204 | 0.0232 | ***** |
| Zenda - Borbet | 0.02381 | 0.54549 | 0.044 | 1 | **NS** |
| Kaas - Chalkewadi | 2.09919 | 0.5406 | 3.883 | 0.00196 | ****** |
| Khamdadevi - Chalkewadi | 2.74276 | 0.53166 | 5.159 | < 0.001 | ******* |
| Thoseghar - Chalkewadi | 2.69679 | 0.5406 | 4.989 | < 0.001 | ******* |
| Zenda - Chalkewadi | 0.98844 | 0.54549 | 1.812 | 0.53975 | **NS** |
| Khamdadevi - Kaas | 0.64357 | 0.5363 | 1.2 | 0.89441 | **NS** |
| Thoseghar - Kaas | 0.5976 | 0.54516 | 1.096 | 0.92945 | **NS** |
| Zenda - Kaas | -1.11075 | 0.55001 | -2.02 | 0.40215 | **NS** |
| Thoseghar - Khamdadevi | -0.04597 | 0.5363 | -0.086 | 1 | **NS** |
| Zenda - Khamdadevi | -1.75433 | 0.54122 | -3.241 | 0.02031 | ***** |
| Zenda - Thoseghar | -1.70836 | 0.55001 | -3.106 | 0.03117 | ***** |

**Appendix S8.** Table representing the distance between Plateaus (Km) in the present study. The minimum and maximum distance between two plateaus are approximately 7 Km and 200 Km respectively.

|  | Kaas | Thoseghar | Chalkewadi | Barki | Borbet | Khamdadevi |
| --- | --- | --- | --- | --- | --- | --- |
| Kaas | 0 | 15.94 | 15.69 | 133.48 | 134.6 | 206.5 |
| Thoseghar | 15.94 | 0 | 7.37 | 120.4 | 118.72 | 191.25 |
| Chalkewadi | 15.69 | 7.37 | 0 | 121.65 | 120.19 | 191.1 |
| Zenda | 89.23 | 74.95 | 73.56 | 79.04 | 46.21 | 118.67 |
| Barki | 133.48 | 120.4 | 121.65 | 0 | 89.95 | 140.8 |
| Borbet | 134.6 | 118.72 | 120.19 | 89.95 | 0 | 72.34 |
| Khamdadevi | 206.5 | 191.25 | 191.1 | 140.8 | 72.34 | 0 |

**Appendix S9:** Table representing the details of the study area.

| **Location** | **Latitude** | **Longitude** | **Altitude** | **Area (Km2)** |
| --- | --- | --- | --- | --- |
| Kaas | 17.72919667 | 73.818125 | 1230 m. (4035 ft) | 3.25 |
| Toseghar | 17.58100167 | 73.90571167 | 1147 m. (3763 ft) | 3 |
| Chalkewadi | 17.58511667 | 73.81795167 | 1145 m. (3757 ft) | 13 to 15 |
| Zenda | 16.92081333 | 73.80121333 | 1022 m. (3353 ft) | 1 to 1.5 |
| Barki | 16.74447222 | 73.84666667 | 978 m. (3209 ft) | 2.5 |
| Borbet | 16.51613333 | 73.89186667 | 974 m. (3196 ft) | 0.23 |
| Khamdadevi | 15.87345167 | 74.03167167 | 844 m. (2769 ft) | 4.5 to 5 |
